# Protein-bound sialic acid in saliva contributes directly to salivary anti-influenza virus activity

**DOI:** 10.1101/2021.03.05.433813

**Authors:** Kaori Kobayashi, Chika Shono, Takuya Mori, Hidefumi Kitazawa, Noriyasu Ota, Yuki Kurebayashi, Takashi Suzuki

## Abstract

The oral cavity is an entrance for respiratory viruses, such as influenza. Recently, saliva has been shown to exert both antimicrobial and antiviral activities. Thus, saliva may be a biological factor that contributes to the prevention of influenza infection. However, the actual salivary anti-influenza A virus (IAV) activity in individuals and its determinant factors are unknown. By assessing individual variations in salivary anti-IAV activity in 92 people using an established new high-throughput system in this study, we found that the anti-IAV activity varied widely between individuals and showed a significant positive correlation with protein-bound sialic acid (BSA) level (ρ=0.473; *p* < 0.001). Furthermore, the anti-IAV activity of saliva with enzymatically reduced BSA content was significantly lower. These results indicate that BSA is a direct regulator of salivary anti-IAV activity and is a determinant of individual differences. Additionally, after comparing the anti-IAV activity across the groups by age, anti-IAV activity in young people (aged 5–19 years) were lower than in adults aged 20–59 years and elderly people aged 60–79 years. Our study suggests that BSA levels in saliva may be important in preventing influenza infection.

## INTRODUCTION

Influenza viruses are some of the most important human pathogens. Seasonal influenza viruses (types A and B) cause epidemics worldwide, with the World Health Organization estimating hundreds of million infections and approximately 250,000– 500,000 deaths per year [1, 2], leading to substantial social and economic burdens worldwide [3]. Several preventive and curative strategies exist against influenza, including vaccination and therapeutic agents, such as neuraminidase inhibitors and adamantanes [4]. However, influenza viruses evolve rapidly, which enables them to escape immunity induced by prior infections or vaccination; moreover, those viruses are transmitted efficiently from human to human via respiratory droplets. Recently, there has been substantial concern about the possibility of a pandemic of a novel influenza strain [5], with reports of an increasing resistance of influenza A viruses to neuraminidase inhibitors and adamantanes [6]. Thus, new approaches are needed to effectively prevent and treat influenza infections.

The oral cavity is one of the entry sites for respiratory viruses. Saliva plays an essential role in maintaining the integrity of human health. It provides lubrication and protection and buffering action and clearance in the oral cavity, maintains tooth integrity, and aids taste and digestion [7]. A previous study indicated that hyposalivation may be a risk factor for acute respiratory infection [8], suggesting that saliva may contribute to the role of innate immunity in the early stages of infection. Saliva induces both antimicrobial and antiviral activities [9]. Additionally, multiple soluble factors in saliva have potent inhibitory activity against influenza A viruses (IAVs), including sialic acid-containing mucin 5 B (MUC5B), salivary scavenger receptor cysteine-rich glycoprotein-340 (gp-340), alpha-2-microglobulin (A2M), and lectins (surfactant protein D [SP-D]) [10-18]. Two main mechanisms for the anti-IAV activity of these factors have been proposed. MUC5B, gp-340, and A2M acts as classic γ inhibitors, mediating the inhibition of IAV by presenting a sialic acid ligand that binds to the viral hemagglutinin (HA), thereby preventing HA attachment to target cells [12-15]. In contrast, SP-D acts as a classic β inhibitor by binding the lectin carbohydrate recognition domain (CRD) to specific oligosaccharides on the head of the viral HA [16-18]. It has been shown that salivary anti-IAV activity is primarily due to sialic-acid-containing proteins rather than lectin [13, 19]. As the first line of defense, salivary antiviral factors are likely to play an important role in protection against respiratory infections. However, since previous studies on salivary anti-IAV activity mentioned above have only been conducted using purifying salivary proteins and small size analyses, the actual salivary anti-IAV activity of each individual, and its determinant factors are unknown. To solve this problem, high-throughput evaluation systems are required to evaluate multiple samples simultaneously. In this study, we constructed a new high-throughput system to evaluate salivary anti-IAV activity using neuraminidase activity derived from influenza virus as an index, and we investigated individual variations in salivary anti-IAV activity and its determinant factors.

## MATERIALS AND METHODS

### Participants

In Study 1 of the individual differences in salivary anti-IAV activity, 92 healthy voluntary participants aged 20–59 years (67 men and 25 women; mean age ± S.D., 36.9 ± 9.9 years) were recruited using snowball sampling. This trial was registered with the University Hospital Medical Information Network (UMIN; http://www.umin.ac.jp/; Registration No. 000024720). In Study 2 of age-related changes in saliva, 240 participants aged 5–79 years (60 young people, 30 boys and 30 girls aged 5–19 years; 120 adults, 60 men and 60 women aged 20–59 years; and 60 elderly people, 30 men and 30 women aged 60–79 years), who registered using a monitor at a contractor (Research and Development, Inc.) were recruited. In these studies, the exclusion criteria were as follows: 1) patients with serious disease; 2) those going to a hospital regularly and under regular pharmaceutical treatment; 3) those who were pregnant or expecting pregnancy; 4) those who were physically unable to complete the study-related questionnaires; 5) those with scratches in the oral cavity; and 6) those considered unsuitable to participate in the study by the study physician. All study protocols were approved by the institutional review board of the Kao Corporation and were performed in accordance with the Declaration of Helsinki. After receiving a full explanation of the study, all participants and their legal guardians provided written informed consent.

### Sample collection and treatment

Saliva samples were collected in a quiet room during the morning, between 09:00 and 10:00, and at least 60 min after eating. Subjects were asked to accumulate saliva in their mouths and spit into a sterile plastic dish at the desired time or indicated time, and saliva samples were collected for 5 min in Study 1 or 10 min in Study 2. The saliva secretion rate (mL/min) was calculated by dividing the amount of saliva collected at the time of collection. Whole saliva samples were centrifuged at 15,000 rpm for 15 min at 4°C. The supernatant was collected and stored at −80°C until use. Frozen saliva samples were thawed and vigorously mixed by vortexing prior to the assay.

### Cells and viral culture

Madin–Darby canine kidney (MDCK) cells were cultured in minimum essential medium (MEM; Sigma-Aldrich Co. LLC, St. Louis, MO) supplemented with 5% (v/v) heat-inactivated fetal bovine serum (FBS; Sigma-Aldrich Co.) and 50 µg/mL gentamicin at 37°C in humidified air containing 5% CO_2_. Influenza A virus strains [A/Puerto Rico/8/34 (H1N1)] and [A/Memphis/1/1971 (H3N2)] were propagated at 37°C using MDCK cells in serum-free medium (SFM; Thermo Fisher Scientific Japan K.K., Kanagawa, Japan) supplemented with 2 µg/mL acetylated trypsin (Sigma-Aldrich Co.) and 50 µg/mL gentamicin.

### Saliva sample preparation

For removal of sialic acid of saliva, 100 µL of saliva sample was mixed with 2 µL of 10 U/µL α2-3,6,8-Neuraminidase (Sialidase) from Arthrobacter ureafaciens (Roche) and reacted at 37°C for 30 min. Heat-treated saliva sample was obtained by keeping 100 µL of saliva sample at 95 °C for 30 min in a dry heat block. These sample was tested in the following assays.

### Anti-IAV activity assays of saliva

MDCK cells (4×10^4^ cells/well) were cultured in a 96-well plate at 37°C for 24 h. The saliva samples were diluted eight-fold in phosphate-buffered saline (PBS). Fifty microliters of each saliva sample was mixed with 50 µL of the virus dilutions at a multiplicity of infection (MOI) of 0.5 in PBS and 100 µL of 2× serum-free medium (SFM) at room temperature, and the 180-µL mixture was immediately applied to the MDCK cells. After incubation at 37°C for 30 min, the MDCK cells were washed with PBS and incubated in 100 μL of SFM at 37°C for 16–18 h for virus multiplication. Thirty microliters of the MDCK cell supernatant was mixed with 20 µL of 0.25 mM 2′-(4-methylumbelliferyl)-α-d-N-acetylneuraminic acid (4MU-Neu5Ac; Funakoshi) and incubated at 37°C for 30 min. The enzyme reaction for 4MU-Neu5Ac was stopped by adding 200 µL of 100 mM sodium carbonate buffer (pH 10.7). Fluorescence from the sialidase reaction of virus NA was quantified using a microplate reader (Infinite 200 multi-mode Reader, Tecan, Männedorf, Switzerland) with excitation and emission wavelengths of 355 nm and 460 nm, respectively, for 4-metylumbelliferone (4MU). The relative sialidase activity in each sample was calculated by setting the sialidase activity to 100% when PBS was used instead of saliva. Finally, anti-IAV activity was defined by deducting the relative sialidase activity from 100%.

### Hemagglutination inhibition (HI) assay

To remove the nonspecific inhibitory activity of saliva-induced hemagglutination, 100-µL saliva samples were mixed with 200 µL of guinea pig erythrocytes and incubated at 4°C for 2 h. The mixture was centrifuged at 2,000 *g* for 10 min, and the supernatant (pretreated saliva sample) was used for the HI assay. The PR8 IAV strain was adjusted to 4 hemagglutinin units (HAU) per 25 µL PBS using a pre-hemagglutinin assay. In 96-well plates, a two-fold serial dilution of the pretreated saliva sample was mixed with 25 µL of 4-HAU PR8 IAV strain, followed by the addition of 50 µL of 0.7% guinea pig red blood cells and left at 4°C for 2 h. The HI titer was calculated as the reciprocal of the highest dilution that produced complete hemagglutination inhibition.

### Sialic acid assay

The concentrations of total sialic acid (TSA) and free sialic acid (FSA) in saliva were measured using a sialic acid assay kit (Bioassay Systems). The concentration of protein-bound sialic acid (BSA) was then calculated by subtracting the concentration of FSA from the concentration of TSA.

### Salivary proteome analysis

For ultrafiltration, 300 µL of saliva sample was mixed with 200 µL of 100 mM Tris-HCl buffer (pH 8.5; Nippon Gene Co., Ltd.) in an Amicon Ultra-0.5 Centrifugal Filter Unit (3 kDa; Merck Millipore Ltd), and the mixture was spun down at 14,000 *g* for 30 min at 4°C. The trapped saliva on the filter was filtered three times using 400 µL of Tris-HCl buffer. Finally, the filter on a new tube was spun down at 1,000 *g* for 2 min at 4°C, and the sample for proteome analysis was collected. The total protein concentration of each sample was determined using the Pierce(™) BCA Protein Assay Kit (Thermo Fisher Scientific Inc.) and the saliva sample was prepared with equal concentrations of proteins from each individual sample. The saliva sample containing 100 µg protein in Tris-HCl buffer (90 µL in total) was mixed with 5 µL of 100 mM DL-dithiothreitol (MP Biomedicals, LLC) for the reduction solution and heated at 56°C for 45 min. The mixture was added to 5 µL of 111 mg/mL iodoacetamide (FUJIFILM Wako Pure Chemical Corporation) and incubated at 37°C for 30 min in shade for alkylation. The mixture was then mixed with 50 µL of 0.1 µg/µL trypsin solution (Promega Corporation) and incubated at 37°C for 16 h. The reaction solution was added to a final concentration of 0.3% (v/v) of 10% trifluoroacetic acid (FUJIFILM Wako Pure Chemical Corporation) to stop the reaction, and then subjected to LC-MS/MS analysis. All reagents were diluted with Tris-HCl buffer. Five µL of the purified peptides were injected into a high-performance liquid chromatography system (ACQUITY UPLC: Waters) connected to a triple-quadrupole mass spectrometer (TSQ Vantage; Thermo Fisher Scientific) with an ion source of electrospray ionization in multiple reaction monitoring (MRM) mode. Chromatographic separation was performed on a Cadenza CD-C18 column (150 × 1 mm, 3 µm; Imtakt) by binary gradient elution at a flow rate of 0.05 mL/min. Eluent A was 0.1% formic acid in water, while acetonitrile containing 0.1% formic acid was used as eluent B. Ion source (ESI) parameters were optimized as follows: spray voltage, 400 V; vaporizer temperature, 100°C; sheath gas pressure, 40 Arb; auxiliary gas pressure, 5 Arb; and capillary temperature, 250°C. The peak areas for each target peptide were calculated using Xcalibur (Quan Browser) software (Thermo Scientific).

### Enzyme-linked immunosorbent assay (ELISA)

The concentrations of secretory IgA (sIgA), defensin alpha 1, neutrophil (DEFa1), and LL-37 in saliva were measured using competitive ELISA kits (sIgA; Immundiagnostik GmbH, DEFa1; CLOUD-CLONE CORP., LL-37; Hycult Biotech Inc.) in accordance with the manufacturer’s instructions.

### Statistical analysis

Results from multiple experiments are expressed as means ± standard deviation (S.D.) and all data were analyzed using IBM SPSS Statistics Version 24 (IBM Japan, Ltd., Tokyo, Japan). Relationships between the two sets of data were analyzed using Spearman’s rank correlation. The statistical significance of differences between groups was determined using one-way analysis of variance, followed by Tukey’s (Fig. 3) or Kruskal-Wallis (Fig. 4) multiple comparisons test. Results with *p* < 0.05 were considered statistically significant.

## RESULTS

### Construction of high-throughput evaluation system for salivary anti-IAV activity

To simultaneously evaluate the anti-IAV activity in many saliva samples, we constructed a new high-throughput evaluation system using neuraminidase (NA) derived from IAV as an index. Fig. 1A shows the schematic of the measurement flow of salivary anti-IAV activity. The measurement consisted of the following steps: 1) mixing the virus and saliva, 2) infection of MDCK cells using the IAV solution and saliva mixture, 3) collecting the supernatant of the cells after 16–18 h of infection, and 4) measurement of sialidase activities of the supernatant for multiplicated virus quantification. The NA activity value when PBS was used instead of saliva was set to 100%, and the relative value was calculated from the NA activity value of saliva samples. The value obtained by subtracting the relative NA activity value from 100 was defined as the anti-IAV activity. A high correlation (ρ=0.916; *p* < 0.001) was confirmed between the NA activity value in the supernatant of infected cells and the number of infected cells stained for NP protein when MDCK cells were infected with IAVs mixed with various saliva samples (Fig. 1B).

**Figure 1.**
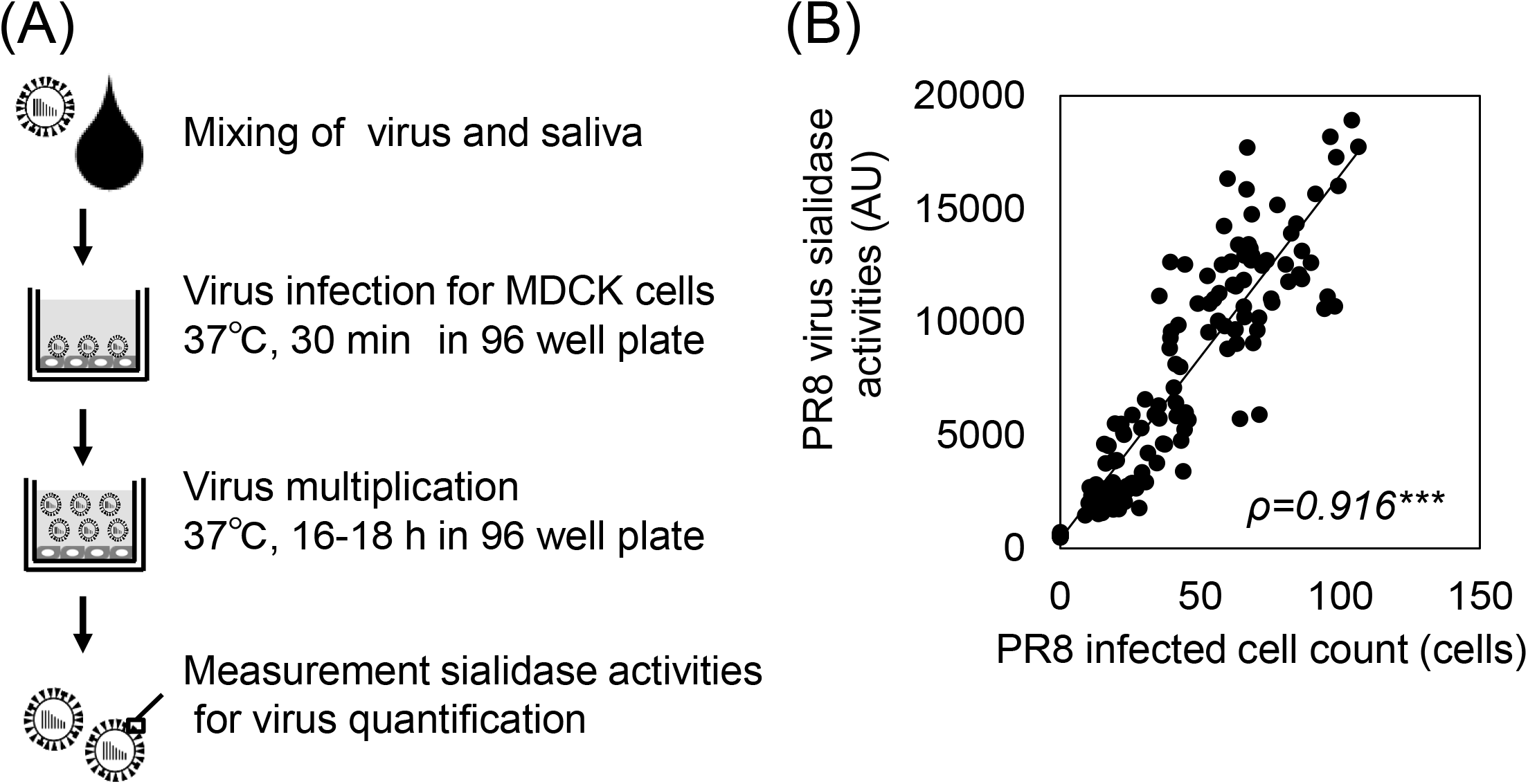
Construction of high-throughput assay of salivary anti-IAV activities. (A) Schematic of the measurement of salivary anti-IAV activities. (B) Correlation between viral sialidase activity and infected cell count. The plot indicates significant moderate positive correlation. ρ, Spearman’s correlation coefficient. ***, *p* < 0.001

### Individual differences in salivary anti-IAV activity

Using the established system, salivary anti-IAV activity against the PR8 IAV strain (H1N1) in 92 individuals was assessed. Anti-IAV activity varied widely between individuals, ranging from 2% to 97% (Fig. 2A). Furthermore, salivary anti-IAV activity against the Memphis IAV strain (H3N2) was significantly positively correlated with anti-IAV activity against the PR8 IAV strain (ρ=0.638; *p* < 0.001) (Fig. 2B). We then performed a correlation analysis of salivary anti-IAV activity against the PR8 IAV strain and the amount of various salivary components measured using proteomic analysis and some kits to explore factors associated with individual differences in salivary anti-IAV activity (Fig. 2C). The results showed the highest positive correlation between salivary anti-IAV activity and BSA levels (ρ=0.473; *p* < 0.001). In addition, the conentration of several proteins, such as AGTN/gp-340, ZG16B, SGP28, and C6orf58, significantly positively correlated with anti-IAV activity.

**Figure 2.**
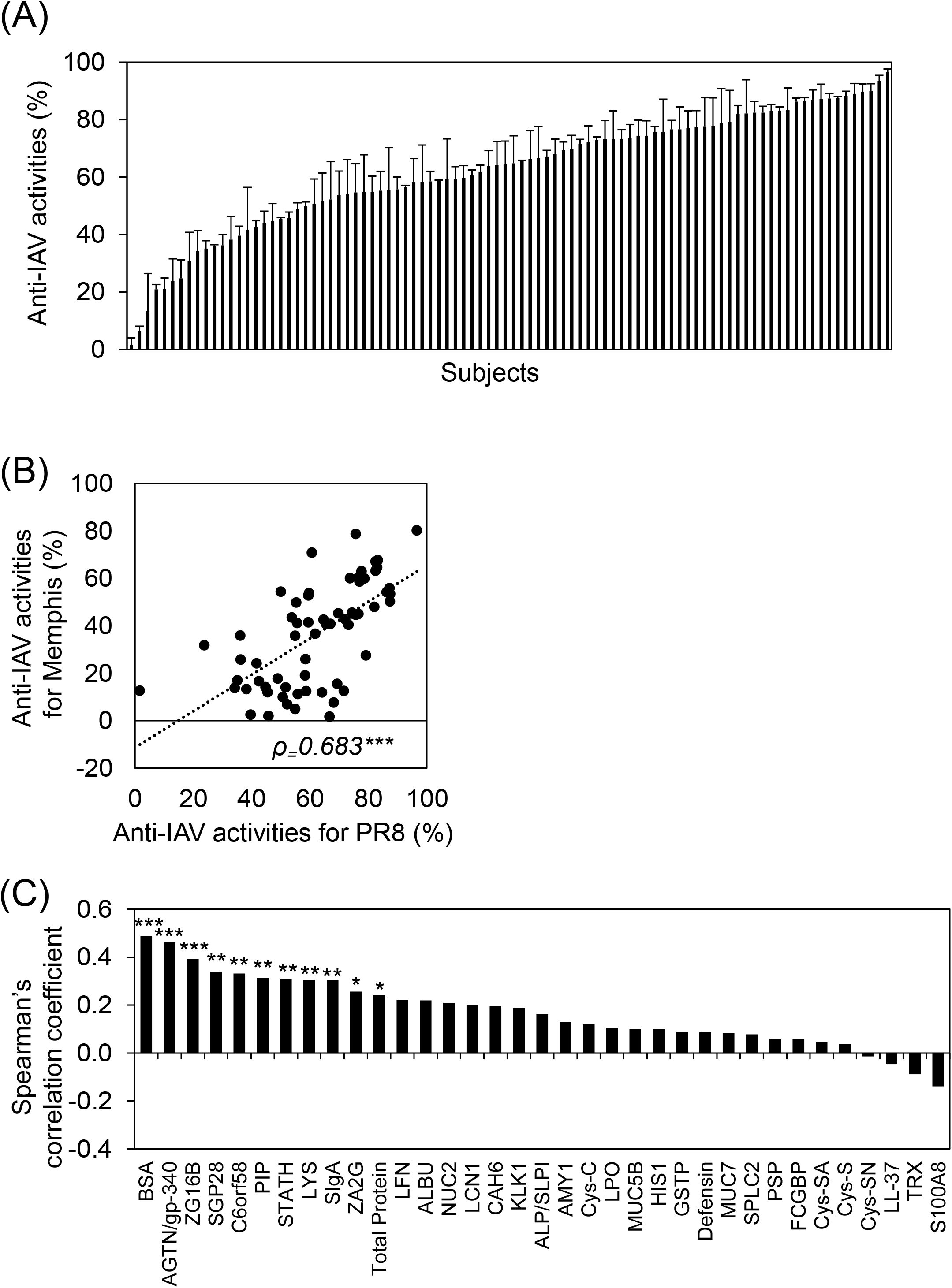
Individual differences of salivary anti-IAV activities. (A) Salivary anti-IAV activities against PR8 IAV strain (n=92). Data are presented as mean ± SD from three independent experiments. A bar graph shows the activity in each subject. (B) Correlation between salivary anti-IAV activity against PR8 and Memphis IAV strains (n=65). The plot shows significant moderate positive correlation. (C) The correlation between salivary anti-IAV activity against the PR8 IAV strain and amount of various salivary components (n=72–85). A bar graph shows Spearman’s correlation coefficient (ρ) in each subject. *, *p* < 0.05; **, *p* < 0.01; ***, *p* < 0.001

### Contribution of BSA to salivary anti-IAV activity

To clarify whether BSA contributes directly to the anti-IAV activity of saliva, we assessed saliva samples that were heat-treated or in which BSA was enzymatically removed. The amount of BSA in the saliva was significantly reduced by α2-3,6,8-sialidase treatment and was not altered by heat treatment (Fig. 3A). While the anti-IAV activity of untreated saliva samples averaged about 50%, the anti-IAV activity of heat-treated saliva samples decreased to an average of 25% (Fig. 3B). In contrast, the anti-IAV activity of sialidase-treated saliva samples decreased to less than 10% on average (Fig. 3B). To further elucidate the contribution of BSA to salivary anti-IAV activity, a hemagglutination inhibition (HI) test was performed. Similar to the anti-IAV activity, large individual differences were observed in HI titers in some saliva samples (Fig. 3C). In addition, individual HI titers highly positively correlated with anti-IAV activity (Fig.3D). Although no effect of heat treatment on HI titer was observed, sialidase treatment significantly decreased the HI titer (Fig. 3E).

**Figure 3.**
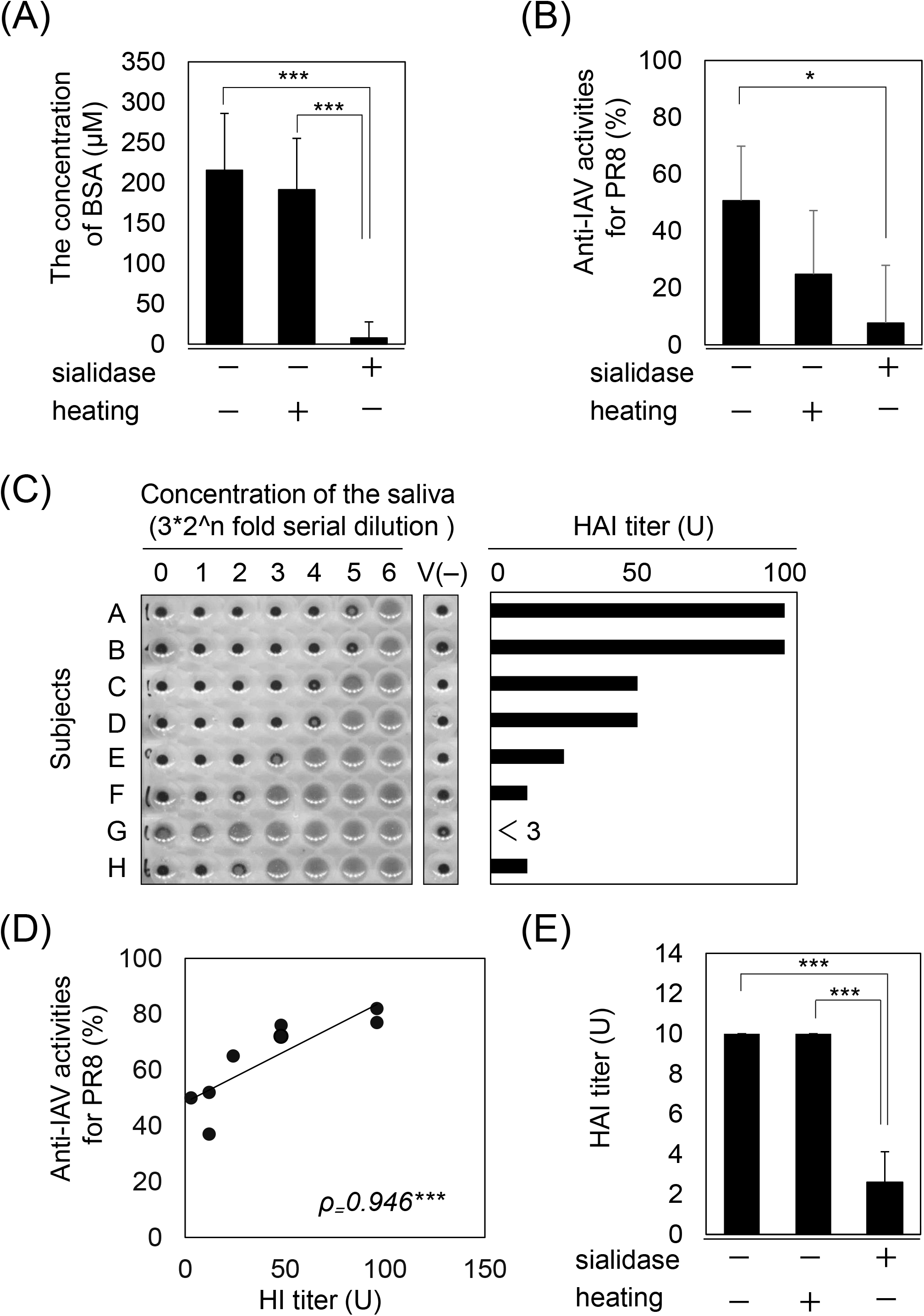
Contribution of protein-bound sialic acid to salivary anti-IAV activity. Effect of α2-3,6,8-sialidase or heat treatment on (A) the concentration of BSA and (B) salivary anti-IAV activity against the PR8 IAV strain. (C) Hemagglutination inhibition assay with saliva (n=8, A-H). The bar graph shows the HI titers of each subject. This experiment was repeated three times with similar results. (D) Correlation between salivary anti-IAV activity against the PR8 IAV strain and the HI titer of saliva. The plot shows a moderately positive correlation. (E) The effect of α2-3,6,8-sialidase and heat treatment on the HAI titer of saliva. Data are presented as mean ± SD for each group. Average plateau values were compared using one-way analysis of variance and Tukey’s test. ρ, Spearman’s correlation coefficient. *, *p* < 0.05; ***, *p* < 0.001.

### Age-related differences in salivary anti-IAV activity

We then compared salivary anti-IAV activity across a wide range of ages (5–79 years divided into three groups, young people (aged 5–19), adults (aged 20–59 years), and elderly people (aged 60–79 years), using the high-throughput evaluation system we constructed. The saliva secretion rate decreased with increasing age and was lowest in elderly people (Fig. 4A). In contrast, the salivary anti-IAV activity was similar in adults and elderly people, and was significantly lower in young people (Fig. 4B). In each age group, the salivary anti-IAV activity significantly positively correlated with salivary BSA level (young; ρ=0.366, *p* < 0.05, middle adults; ρ=0.589, *p*-value < 0.001, elderly; ρ=608, *p*-value < 0.001, total; ρ=0.511, *p*-value < 0.001) (Fig. 4C).

**Figure 4.**
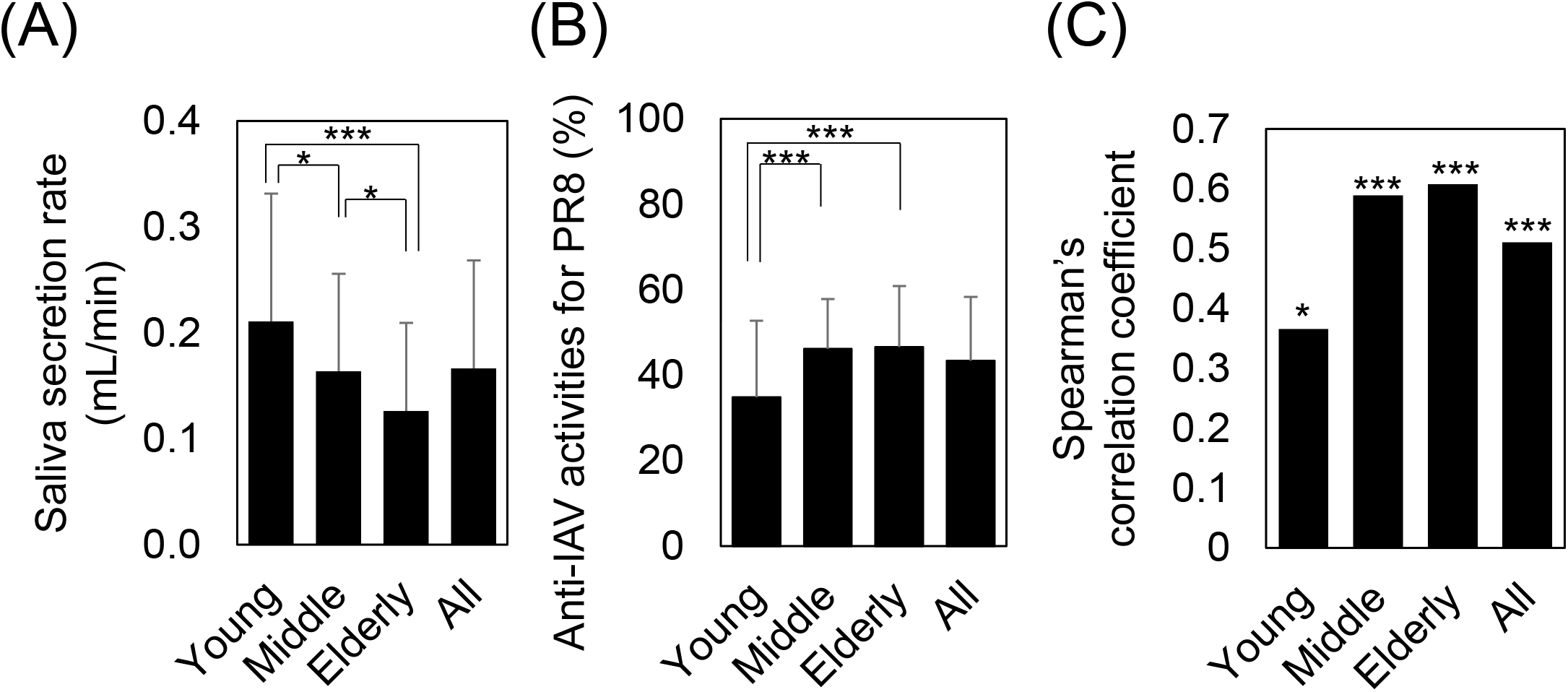
Age-related differences in salivary anti-IAV activity. Comparison of (A) the total saliva volume (mL/min), (B) salivary anti-IAV activit against the PR8 IAV strain, and (C) correlation between salivary anti-IAV activit against the PR8 IAV strain and the concentration of BSA in young people (30 boys and 30 girls aged 5–19 years), adults (60 men and 60 women aged 20–59 years), and 60 elderly people (30 men and 30 women aged 60–79 years). Data are presented as mean ± SD of each group. Average plateau values were compared using one-way analysis of variance and the Kruskal-Wallis test. *, *p* < 0.05; ***, *p* < 0.001.

## DISCUSSION

In this study, we constructed a high-throughput system to evaluate salivary anti-IAV activity. Using this system, we found that there are large individual differences in salivary anti-IAV activity, and that the amount of BSA in saliva is important for these individual differences.

In general, the main method of measuring the viral infection titer is to count the number of infected cells by staining with antibodies against the virus and clumps of cellular degeneration associated with viral infection. While these methods are suitable for measuring the virus titer, it is difficult to evaluate multiple samples because this approach requires a great deal of time and labor because of the need for operations, such as multi-step dilutions and infected cells or colony couting. In our constructed system, the neuraminidase activity derived from the propagated influenza virus in the culture medium was measured using a multi-plate reader, which eliminates the operations described above and is suitable for the analysis of multiple samples. A significant positive correlation was observed between salivary anti-IAV activity against the PR8 (H1N1) and Memphis (H3N2) strains (Fig. 2B), indicating that individual differences in salivary anti-IAV activity are similar across influenza virus strains. It is expected that this method can be used in the evaluation of many influenza virus strains in the future.

Using the constructed system, it was clarified that the individual variations in salivary anti-IAV activity most positively correlated with the amount of BSA (Fig. 2C). Moreover, the HI titers of saliva were significantly reduced by treatment with α2-3,6,8-sialidase (Fig. 3B). Some studies have reported that saliva contains soluble factors that can exert anti-influenza activity, such as sialic acid binding protein and lectins. It has been reported previously that several purified salivary proteins, such as gp-340 and A2M, can inhibit the hemagglutination of erythrocytes by presenting a sialic acid ligand for the viral HA [12-15]. These reports are consistent with our results, and indicate that the total amount of sialic acid possessed by these proteins determines the anti-IAV activity of saliva. In other words, the amount of BSA in the saliva is expected to be an indicator of frontline oral defenses against influenza viruses.

MUC5B has potent inhibitory activity against IAV. However, our data showed that salivary MUC5B levels weakly correlated with the anti-IAV activity of saliva. White et al. reported that the sialic acid of MUC5B is more easily cleaved than that of gp-340, the levels of which highly correlated with anti-IAV activity. Our results suggest that the sialic acid of MUC5B is cleaved by sialidase in the oral cavity [20-22], and the anti-IAV activity of MUC5B in saliva may be reduced. In considering individual variations in the anti-IAV activity of saliva, it is necessary to consider oral sialidase activity. In this study, SGP28, C6orf58, and PIP also strongly correlated with the anti-IAV activity of saliva. These molecules are known to have sugar chains, but the presence of sialic acid is not clear [23-25]. They may also contribute to the anti-IAV activity of saliva via sialic acid, which is less susceptible to cleavage by sialidase. It is known that sialic acids are structurally diverse and that O-acetylated sialic acids are resistant to bacterial sialidase [26]. In the future, it will be necessary to clarify the presence or absence of sialic acid in these molecules, including their structural features.

Interestingly, the anti-IAV activity of saliva was reduced following excessive sialidase treatment, but was not completely lost (Fig. 3A, 3 B). Moreover, salivary anti-IAV activity also decreased following heating, which did not affect the concentration of sialic acid. Therefore, it is possible that sialic acid-independent factors are also involved in anti-IAV activity. One of the possible factors is the action of lectin-like proteins, such as ZG16B, which have shown a high correlation with anti-IAV activity [27, 28]. Lectin (carbohydrate-binding proteins), including conglutinin, SP-D, MBL, and SAP, bind selectively to specific carbohydrate structures (mannose-over galactose-type sugars) located at the head of the influenza HA of susceptible strains, thereby blocking the ability of HA to bind to sialylated cell-surface receptors [16, 17]. Furthermore, lectin-like proteins may also inhibit the fusion of the viral membrane with the host cell by binding to sialic acid on the surface of the host cell [18]. In our evaluation system, since host cells were exposed to a mixture of saliva and virus for 30 min, the action of lectin-like proteins in saliva that inhibit the contact between host cells and virus would also contribute to the anti-IAV activity of saliva. On the other hand, in our evaluation system, the activity of molecules that inhibit the later mechanisms of virus propagation, such as replication and release, is not reflected in the anti-IAV activity. For this reason, defensin and lactofferin, which are known to inhibit replication of IAV [12, 29-33], showed only a weak correlation with anti-IAV activity in our study.

The current thinking regarding influenza infection morbidity is that the risk factors are related to age, higher rates of infection in school-aged children relative to adults, and lower rates in the elderly. Children spend a great deal of time in communities where daily contact with other people is extensive; for example, in schools, playgroups, and daycare centers, and it is assumed that close contact favors infection [34, 35]. The results of evaluating differences in salivary properties by age (Fig. 4) showed that salivary volume, which has been reported as a predictive risk factor for influenza transmission [8], was higher in the younger age group. On the other hand, the anti-IAV activity of saliva in the young group was significantly lower than that in the middle and elderly groups. These results suggest that low salivary anti-IAV activity may also be associated with a high rate of IAV transmission in school-aged children. In addition, salivary anti-IAV activity was highly correlated with sialic acid in each age group, indicating that the determinants of anti-IAV activity were not as sensitive to age. Thus, qualitative control of saliva in young people may be effective in preventing IAV infection. Prospective intervention trials with techniques that provide qualitative control of saliva are needed to gain insight into the contribution of salivary anti-IAV activity to IAV infection in young people.

Our study revealed that there are large individual differences in the anti-IAV activity of saliva, which mainly depends on the amount of BSA. BSA directly contributes to salivary anti-IAV, which may contribute to influenza morbidity in young people. In conclusion, the understanding of individual differences in salivary anti-IAV activity may provide new insights into creating effective methods to prevent influenza infection.

## Acknowledgments

We would like to thank Editage (www.editage.com) for English language editing.

## Author contributions

T.M. and N.O. designed the project; K.K., C.S., and H.K. performed the research; K.K., C.S., H.K., T.M., Y.K., and T.S. analyzed the data; K.K., C.S., and T.M. wrote the paper. T.M., N.O., Y.K., and T.S. edited the manuscript.

